# Morphological and genetic analysis of inflorescence and flower development in hemp (*Cannabis sativa* L.)

**DOI:** 10.1101/2024.01.25.577276

**Authors:** Jiaqi Shi, Susanne Schilling, Rainer Melzer

## Abstract

Hemp (*Cannabis sativa*) is one of the oldest cultivated crops, and its myriad of uses have fascinated humans for millennia. In contemporary agriculture, the seeds are used for high-quality oil, human food, and nutritional supplements. From female inflorescences, secondary compounds (cannabinoids, terpenes, and flavonoids) are isolated that are implicated to have a wide range of medicinal applications. These elements provide hemp with significant market and research value and emphasise the need to thoroughly understand reproductive development in hemp.

Here, we present a detailed morphological and molecular analysis of hemp inflorescence and flower development of the cultivar ‘FINOLA’. Hemp is unusual among flowering plants as it is dioecious, i.e. develops male and female plants. We define eight landmark events in male and female flower development and show that developmental differences between male and female plants extend beyond floral morphology, additionally comprising inflorescence structure as well as flowering time. Further, we demonstrate that the time of activation of key reproductive developmental regulators is significantly different in male and female hemp plants. Our comparison of male and female hemp plants shows that developmental pathways diverge very early, already at the two-leaf stage, laying a basis for further exploration into the genetics of sex determination of *C. sativa*.

## Introduction

*Cannabis sativa*, a member of the family Cannabaceae, holds great potential as a high-value crop. Hemp seeds are rich in Omega-3 fats and make for a beneficial dietary supplement for humans (Leizer et al., 2000). Additionally, the medicinal compound CBD has been reported to treat epilepsy effectively (Silvestro et al., 2019), and ongoing research is exploring its potential benefits for treating inflammation, depression, and cancer (Fraguas-Sánchez and Torres-Suárez, 2018). Furthermore, hemp can be utilised for renewable bio-based plastics, textiles and construction materials (Finnan and Burke, 2013). In addition to its economic value, hemp is considered an eco-friendly and sustainable crop due to its short growth period, low water requirements, and high carbon-sequestration potential (Finnan and Styles, 2013).

For many of the aforementioned applications, the hemp flower plays a central role. For example, the production of pharmacologically active compounds primarily occurs in female hemp flowers (Spitzer-Rimon et al., 2019). CBD is mainly extracted from the trichomes of female hemp inflorescences (Aizpurua-Olaizola et al., 2016). Therefore, it is essential to better understand the reproductive development of hemp.

Hemp is a primarily dioecious plant, meaning that it has separate male and female individuals. This is a unique characteristic, as only 6% of plants exhibit this trait (Feng et al., 2020). However, with the exception of the well-known mature flower structure (Leme et al., 2020; Spitzer-Rimon et al., 2019), little is known about the dynamic processes of plant growth, development, and flowering in both sexes, including inflorescence development and the stages of flower development.

Moreover, knowledge about the genetic mechanism of hemp floral development is currently limited. It is known that the induction of flowering is controlled by specific genes that act as switches, signalling the transition from vegetative to reproductive organ development. These genes have been extensively studied in *Arabidopsis thaliana* (Arabidopsis) during flower development (Ryan et al., 2015). Previous studies have shown that *SUPRESSOR OF OVEREXPRESSION OF CONSTANS1* (*SOC1*) plays a crucial role in promoting floral meristem identity by activating other flowering genes in Arabidopsis (Liu et al., 2009). Additionally, *LEAFY* (*LFY*) is a key gene involved in floral meristem identity and flowering time. It is an important factor in the transition from the vegetative to the reproductive phase (Blazquez et al., 1997). *AGAMOUS* (*AG*) is a key regulator in the development of stamens and carpels. Its expression is typically highest during the late flowering stages (Okamuro et al., 1993, Ó’Maoiléidigh et al., 2013). It is currently unclear how homologs of those key regulators of flower development are expressed in hemp and whether there are expression differences in male vs female flowers.

Here, we provide a detailed morphological and molecular analysis of landmark events in hemp reproductive development. Since hemp is a dioecious plant with both male and female flowers, our study seeks to comprehensively understand both male and female flower development processes. We show that male and female development diverge already at a very early time point and that male and female plants do not only differ in flower morphology but also in inflorescence architecture and flowering time, indicating a profound difference in the gene regulatory networks governing male and female plant development.

## Materials and Methods

### Plant cultivation and phenotyping

The seeds of *Cannabis sativa* photoperiod-insensitive cultivar ‘FINOLA’ was germinated in a mixture of 1 part perlite, 1 part vermiculite, and 2 parts compost (John Innes No. 2). After four days of germination in darkness, seedlings with a first true leaf of approximately 1 cm in length were transferred to individual pots. Following germination in darkness, seedlings were cultivated under natural light conditions (Dublin, Ireland from June to September 2021) or under artificial light (red– blue LEDs, light intensity approximately 500 μmol/sec at approximately 15 cm distance from the light source; HORTILED MULTI, 44 W; HORTILUX, https://www.hortilux.com/) in long day conditions 16 h of light and 8h of darkness (Schilling et al., 2023). Phenotyping was conducted during the whole growing period. The external phenological growth stages and flower development were systematically characterised and photographed using a digital camera (Panasonic Lumix DC-G9, Japan). The plants were monitored daily after being transferred to individual pots. Two angles were used for observation: side profiles to track the number of leaf pairs and top view to record changes in the shoot apical meristem. Developing inflorescences and flowers were photographed using a stereomicroscope (LEICA M165FC, LEICA V4.1 software, Germany).

### Light microscopy

Plant samples (apical buds) were taken at different stages which were selected from 64 plants in the growth room and immediately fixed in 4% Formaldehyde Alcohol Acetic Acid (FAA) overnight at 4 °C. Subsequently, tissues were dehydrated using an ethanol series of 50%, 70%, 90% and 100%, followed by an ethanol - Neo-clear (Millipore, Germany) series, transitioning the sample into 100% Neo-clear. Finally, the Neo-clear was replaced with paraffin wax (Sigma-Aldrich, America). Sections of 5–8 µm thickness were cut using a microtome (microTec D-69190 Walldorf, Germany), mounted on glass slides and stained using toluidine blue 0.5%. Slides were imaged using a light microscope (Leica DM3000, Germany).

### RNA extraction, purification and cDNA synthesis

Plant tissue (apical structures, including newly emerged leaves up to 10 mm) was harversted during the different vegetative stages and reproductive stages which were selected from 233 plants in the greenhouse, instantly frozen in liquid nitrogen and stored at -80°C. RNA isolation was performed using the RNeasy® plant mini kit (Qiagen, Germany) according to the manufacturer’s instructions for each plant tissue separately.

Isolated RNA quality and quantity were measured using a NanoDrop ND-1000 spectrophotometer (Thermo Scientific, America). The RNA integrity was determined by running a 1% agarose gel. Subsequently, the DNase digestion for removing the residual DNA was carried out using the DNase I, RNase free kit (Thermo Fisher Scientific, America) following the manufacturer’s instructions cDNA synthesis was performed according to the manufacturer’s instructions using the Invitrogen SuperScript IV kit (Thermo Fisher Scientific, America). To ensure equal quantities of RNA in each sample, the required amount of RNA for cDNA synthesis was calculated based on the RNA concentration obtained from the NanoDrop ND-1000 spectrophotometer (Thermo Fisher Scientific, America). Negative controls were created for each template RNA sample by using 1 µl of nuclease-free water instead of reverse transcriptase. These negative controls are referred to as the no-RT controls for subsequent qPCR analysis.

### Primer design and gene-expression analysis by RT-qPCR

The *C. sativa* orthologs of *Ubiquitin (UBQ)* and *Protein phosphatase 2A (PP2A)* were selected as reference genes (Guo et al., 2018). Three candidate genes, *CsSOC1*, *CsLFY*, and *CsAG* from *C. sativa* were analysed. The gene structure was inferred from the prediction available at NCBI (GeneBank ID: 115706939, GeneBank ID:115695615, GeneBank ID: 115697576, respectively). *CsAG* has been termed *CsMADS1* and *CsSOC1* has been termed *CsMADS18* elsewhere (Wan et al., 2021). All designed primers sequences for RT-qPCR (Quantitative reverse transcription polymerase chain reaction) were pre-analyzed by OligoAnalyzer (https://eu.idtdna.com/calc/analyzer) (Integrated DNA Technologies, America) (Supplementary Table S1). Primers were synthesised by Integrated DNA Technologies. After being rehydrated in nuclease-free water to a storage concentration of 50 µM, the primers were diluted to a working solution of 10 µM. The efficiency of each primer was assessed by generating a standard curve through serial dilution, and it was observed that the primer efficiency was around 110% for each primer pair.

To study the candidate gene expression, Fast SYBR Green Master Mix (Thermo Fisher Scientific, Country) was used on the ViiA 7 Real-Time PCR System (Thermo Fisher Scientific, America). Data was analysed with QuantStudio 6 and 7 Pro Real-Time PCR Systems Software (Thermo Fisher Scientific, America). For RT-qPCR, a 10 μl reaction mixture included 5 μl 2x Fast SYBR ™ Green Master Mix, 0.2 μl each of specific forward and reverse primers, 0.6 μl DEPC water, and 4 μl cDNA (0.25ng/μl) or nuclease-free water (NTC). The amplification program consisted an initial denaturation at 95L for 30s, followed by 40 cycles of 95L for 10s and 60L for 30s. Melting curve analysis involved heating to 95°C for 15s, cooling to 60°C for 1 min, and gradually increasing the temperature to 95°C at 0.05L/s, followed by a final step at 95L for 15s. Successful target product amplification was confirmed by observing a distinct and singular peak in each melting curve Each experiment was conducted with three biological replicates and four technical replicates.

## Results

### Male and female hemp plants experience different lengths of vegetative growth

To establish landmark events during *C. sativa* development the photoperiod-insensitive dioecious hemp cultivar ‘FINOLA’ was grown under long day conditions under artificial light (Schilling et al., 2023). After germination and emergence of the two cotyledons, true leaves arose in pairs, with individual leaves opposite to each other and successive pairs of leaves arranged at approximately 90 degrees to each other (Figure 1). We used the number of true leaf pairs to indicate the growth stage of the plant: 1st to 9th true leaf stage (L1 to L9).

**Figure 1.**
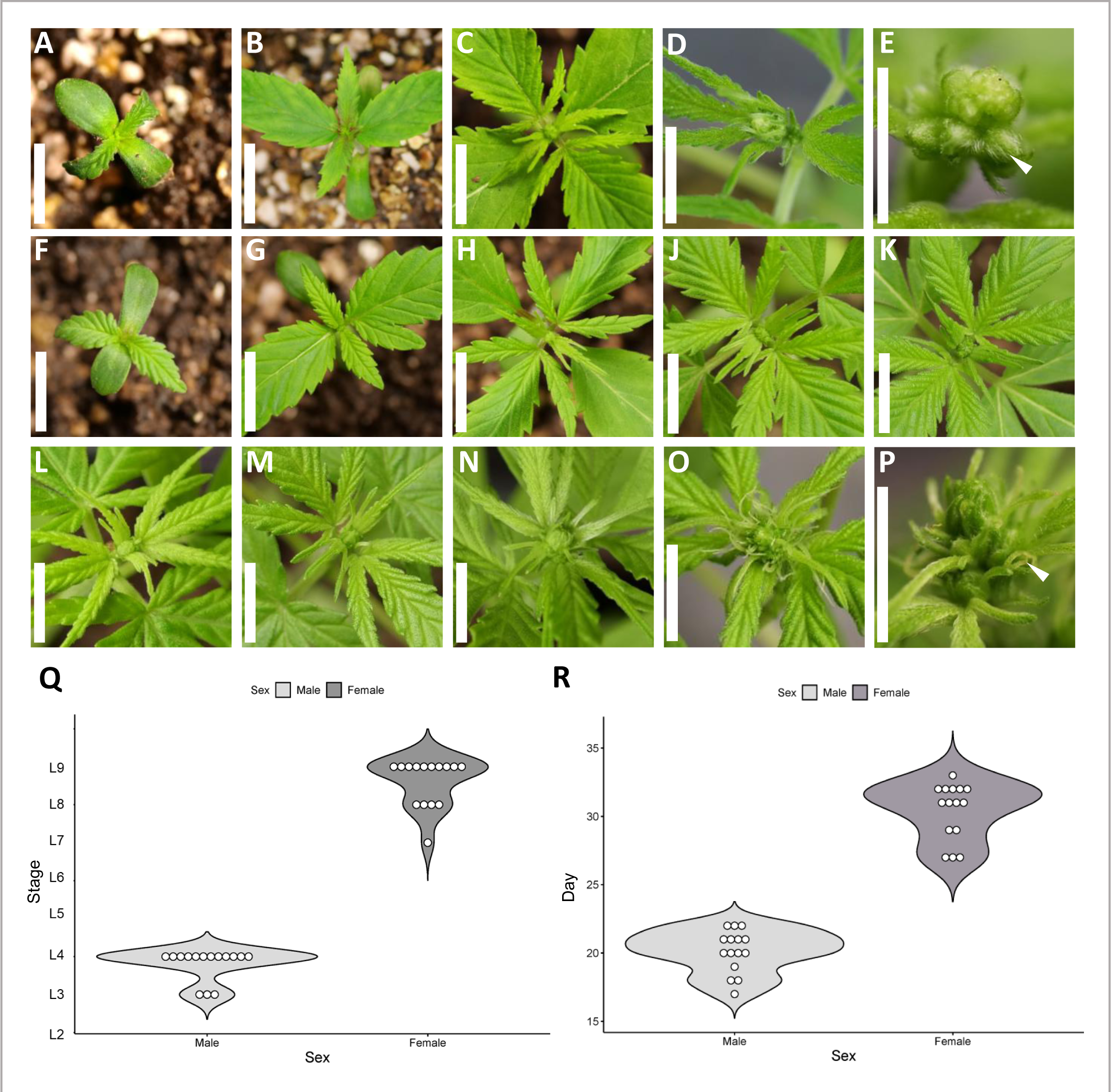
Stages of male and female hemp plants transitioning from vegetative to reproductive development. Male hemp plants of the ‘FINOLA’ cultivar at stage L1 possess cotyledons and an emerging first true leaf pair (A). Male plants at stage L2 exhibit two true leaf pairs (B). At stage L3 (C) the shape of the apical structure of male plants is notable. Male plants at stage L4 (D) have a macroscopically visible apical inflorescence (E), and a single male flower bud pointed by the white arrowhead (E). Female hemp plants of the ‘FINOLA’ cultivar at stage L1 possess cotyledons and an emerging first true leaf pair (F). Female plants at stage L2 have two true leaf pairs (G), at stage L3 showed third true leaf pairs (H) at stage L4 showed fourth true leaf pairs (J), at stage L5 showed fifth true leaf pairs (K), stage L6 showed sixth true leaf pairs (L) and stage L7 showed seventh true leaf pairs (M) all with no visible sign of reproductive development. At stage L8 (N) apical structure in female plants becomes more obvious. Female plants start visibly flowering at stage L9 (O) and the stigma observed from the inflorescence is pointed by the white arrowhead (P). Developmental stage (Q) and days after sowing (R) at which male and female hemp plants start flowering. The scale bar in A to P is 10mm.

In the early stages of vegetative development (L1 and L2), male and female plants were morphologically indistinguishable (Figure 1A, B, F, G). At stage L3, male plants appeared to be vegetative macroscopically, with no distinctive male flowers but a characteristic rounded apical structure (Figure 1C). At stage L4, single flower buds were visible in male plants (Figure 1D, E), indicating transitioning to the reproductive phase.

Compared to male plants, female plants underwent a longer period of vegetative development and continued to produce more true leaves before flowering. Female plants exhibited vegetative characteristics during stages L3 to L7 (Figure 1H to M). At stage L8, the size of the female apical structure obviously increased (Figure 1N). Finally, in stage L9, stigmata were observed in female individuals, showing the female hemp plants started flowering (Figure 1O, P).

The developmental stage as measured by the number of true leaf pairs defined the start of flowering time consistently at L3 and L4 for male plants and L7 to L9 for female plants (Figure 1Q). A greater variability was observed when measuring flowering time in days after sowing (Figure 1R).

### The inflorescence structure differs between male and female hemp plants

As development progressed in male hemp plants of the cultivar ‘FINOLA’, an increase in the number of floral buds was observed (Figure 2A to D). In the male inflorescence, flowers develop sequentially, where more mature flowers are situated at the lower end of the inflorescence (Figure 2E). To gain a detailed understanding of hemp inflorescence development, we conducted longitudinal sections and microscopic analyses. The results showed a large number of developing flowers and floral meristems in the developing inflorescence at growth stage L4. The flowers were at varying stages of development. (Figure 2F). At stage L4 development is rapid, with the number of flowers quickly increasing within the same stage (Figure 2F, G).

**Figure 2.**
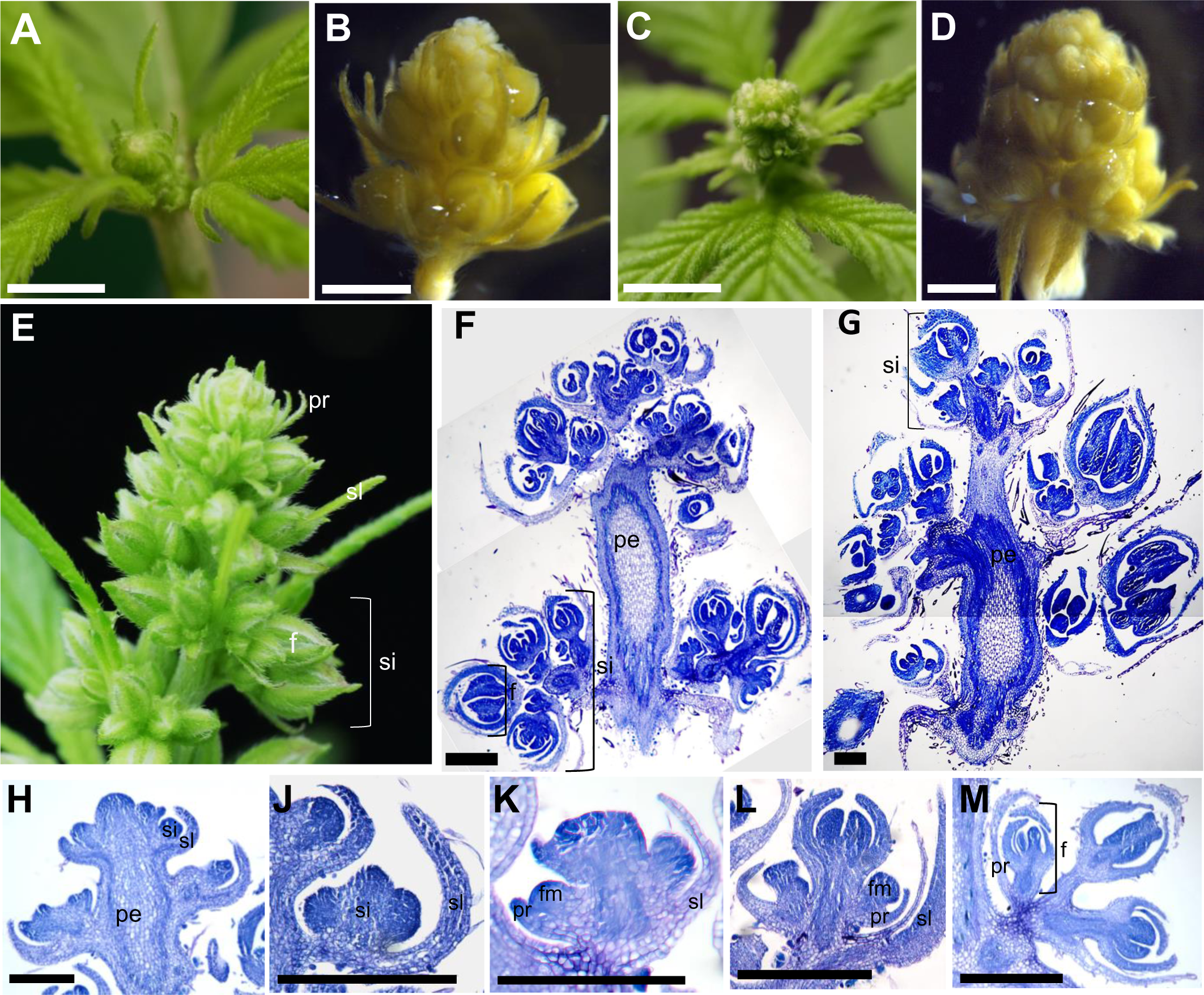
Development and structure of apical inflorescence in male hemp plants. Apical inflorescence at the stage L4 when individual male hemp flowers are first visible macroscopically are shown *in situ* (A) and fixed in ethanol (B). A developmentally more advanced male stage L4 apical inflorescence shown *in situ* (C) and fixed in ethanol (D). A fully developed male inflorescence is composed of multiple secondary inflorescences (E). Longitudinal sections of apical inflorescences at early stage L4 (F) and late stage L4 show flowers at different developmental stages (G). Longitudinal sections show development of the secondary inflorescence from inflorescence meristem (H, J). Floral meristem developed on the secondary inflorescence and embraced by a prophyll (K). Floral meristem (L) and flower (M) are depicted. f. Flower; fm. Floral meristem; pe. Peduncle; pr. Prophyll; si. Secondary inflorescence or secondary inflorescence meristem; sl. Subtending leaf. Scale bar: (A, C) 5 mm; Scale bar: (B, D) 1mm; (F to M) 200 µm.

Male flowers in the inflorescence formed multiple clusters, indicating the presence of secondary inflorescences in the male hemp plant (Figures 2E to G). The secondary inflorescences initiated from the flanks of the apical meristem, and each secondary inflorescence grew from the axil of a subtending leaf (Figure 2H, J). As development proceeded, the floral meristem arose from the secondary inflorescence meristem, with the prophyll or bracteole, a leaf-like structure, extending under each floral meristem (Figure 2 K, L, M) (McMaster and Moragues, 2019).

Similar to male plants, apical inflorescences of female hemp plants gradually developed more flowers after they entered the reproductive phase (Figure 3A to E). As in the male inflorescence, flowers developed acropetally in the female inflorescence, with more mature female flowers located at the bottom of the inflorescence (Figure 3F). Meanwhile, the variation in the degree of development among the flowers was significant, with the flowers at the base of the inflorescence developing floral organs like the ovary while the flowers at the top were still in the floral meristem stage (Figure 3F). In contrast to the structure of the inflorescence in male hemp plants, no clusters of flowers but single floral meristems developed from the flanks of the apical meristem in female inflorescences (Figure 3F, G). Female inflorescences showed a regular pattern of oppositely arranged floral meristems, with each floral meristem enclosed and surrounded by a bracteole (Figure 3F, G). Conceptually, female inflorescences differed in architecture from male inflorescences in the hemp cultivar ‘FINOLA’ under the given light conditions. The female inflorescence can be interpreted as spike-like, while we consider the male inflorescence to be a panicle (Figure 4).

**Figure 3.**
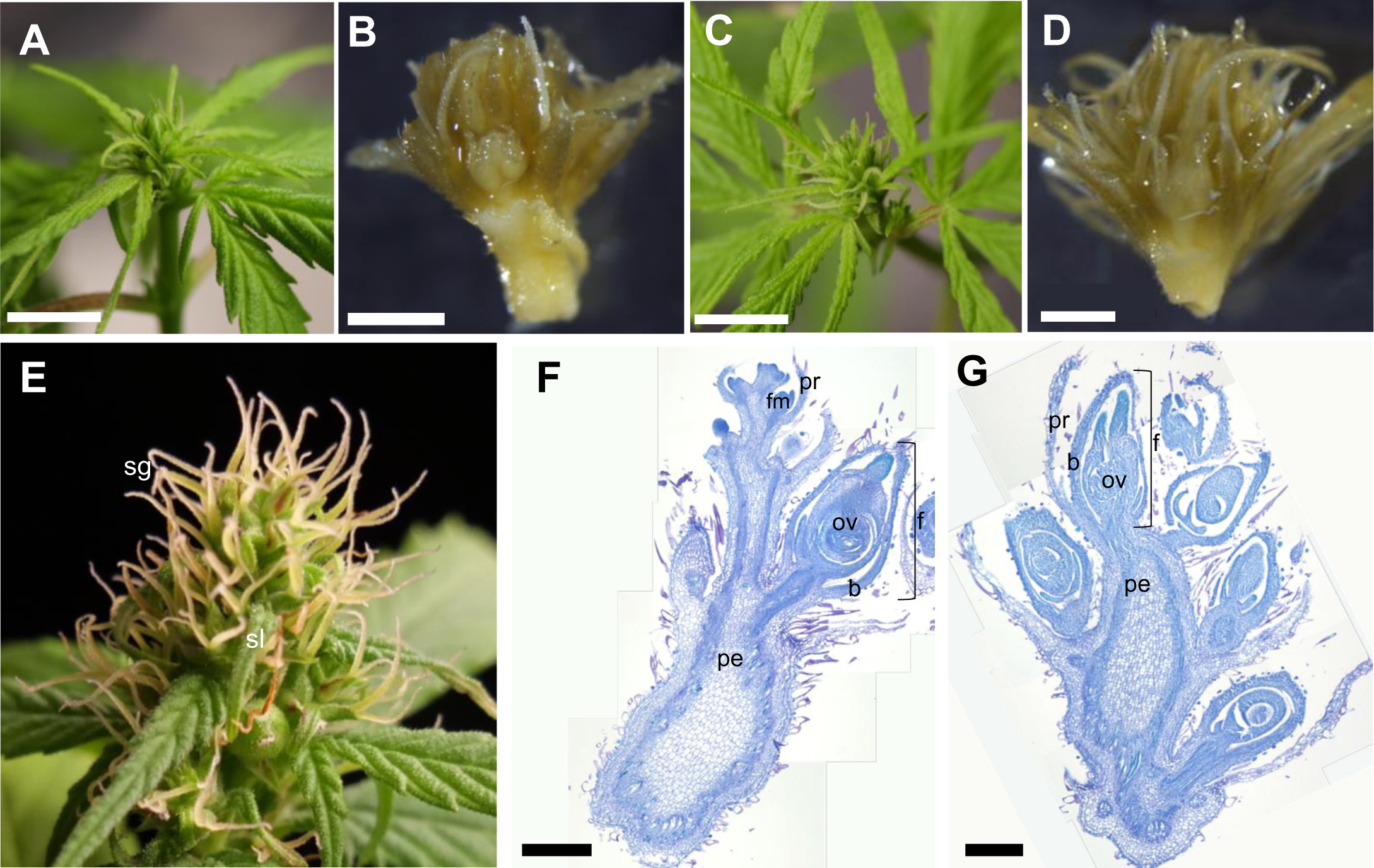
Development and structure of apical inflorescences in female hemp plants. Apical inflorescence at stage L9 when individual female hemp flowers are first visible macroscopically are shown *in situ* (A) and fixed in ethanol (B). A developmentally more advanced female stage L9 apical inflorescence shown *in situ* (C) and fixed in ethanol (D). A fully developed female plant with apical and axillary inflorescences (E). Longitudinal section of female apical inflorescence at early stage L9 (F) and late stage L9 (G). b. Bract; pe. Peduncle; pr. Prophyll; f. Flower; fm. Floral meristem; ov. Ovary; sg. Stigma; sl. Subtending leaf. Scale bar: (A, C) 5 mm; Scale bar: (B, D) 1 mm; (F, G) 200 µm.

**Figure 4.**
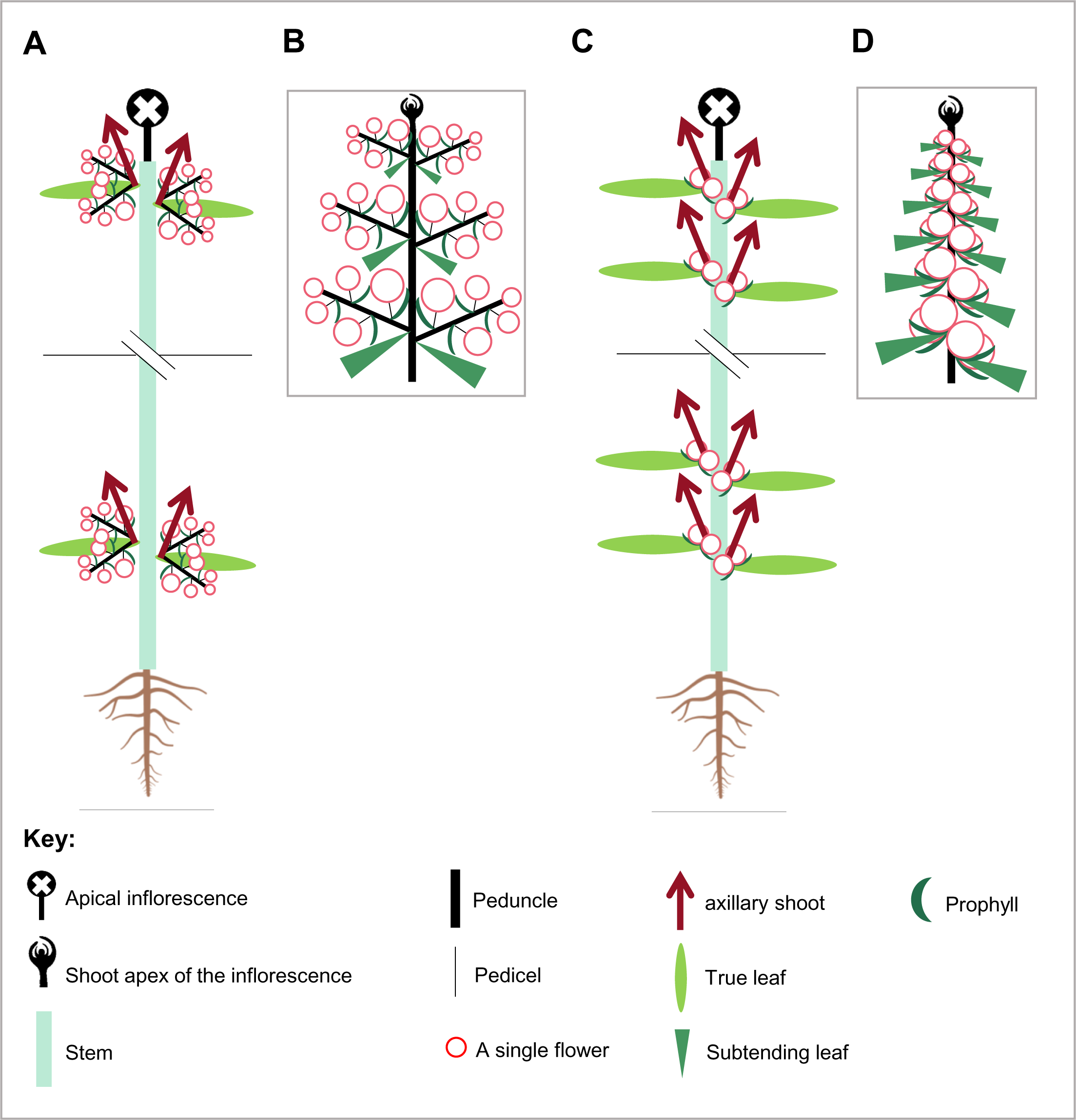
Schematic visualisation of mature hemp plant and inflorescence architecture. Mature male flowering hemp plants (A) and apical inflorescences of the male hemp plant (B). Mature flowering female hemp plants (C) and apical inflorescence of the female hemp plant (D).

### Landmark stages in male and female flower development

Beyond a characterization of the vegetative to reproductive transition in hemp, we aimed to identify landmark events during flower development in hemp (Table 1). The stages of flower development were based on those described previously for Arabidopsis (Smyth et al., 1990).

**Table 1.**
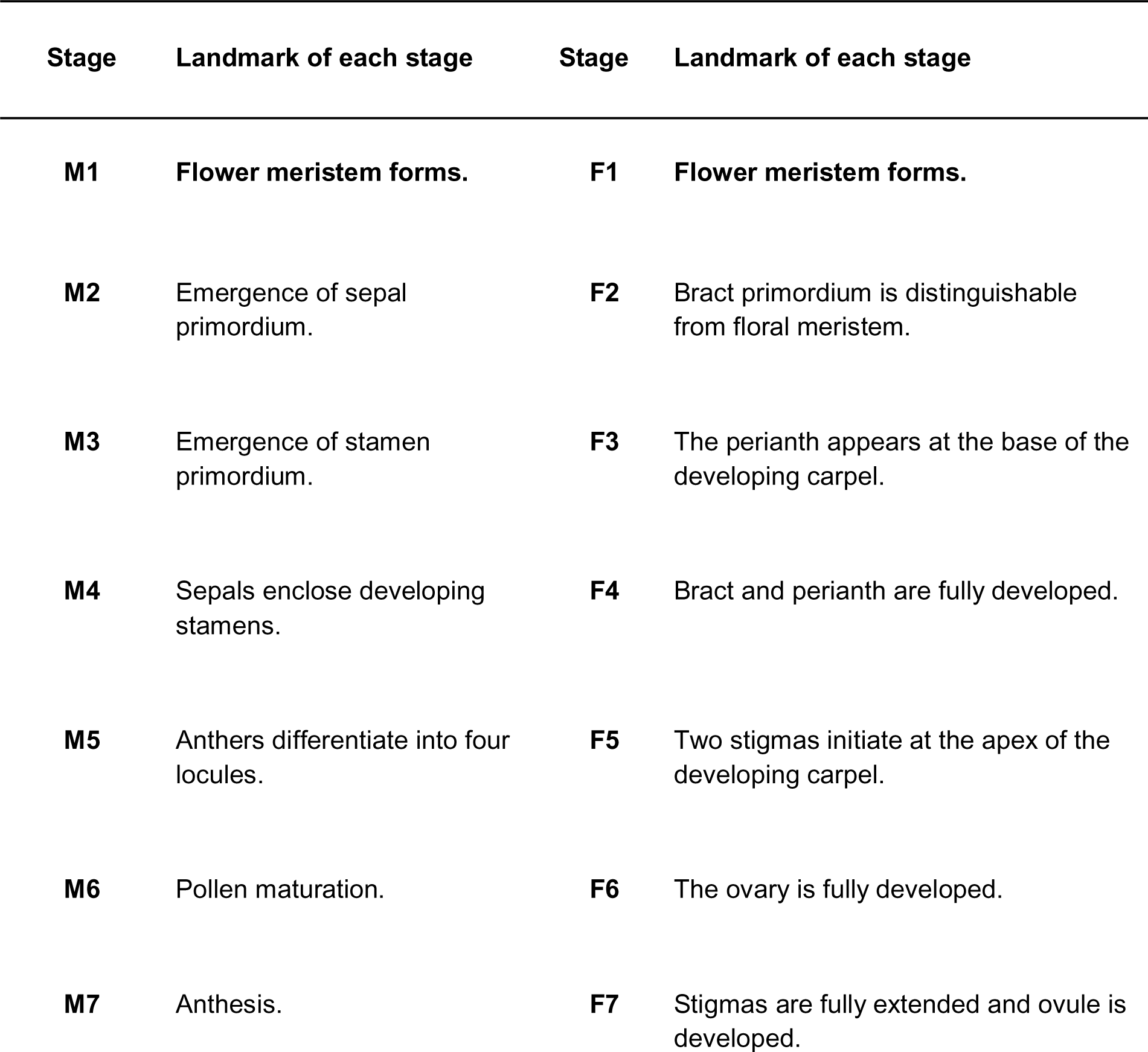
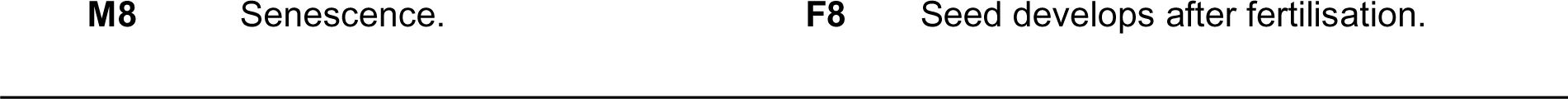
Stages and landmarks of flower development of male and female hemp flowers.

Flower development of male flowers (stage M1) starts with the formation of floral meristem (Figure 5A). At stage M2, sepal primordia arise from the floral meristem (Figure 5B). The stamen primordia appear at stage M3 (Figure 5C). Subsequently, male flower organs grow, and the sepals enclose the developing stamen in stage M4 (Figure 5D). In stage M5 sepals and stamens continue to extend and develop. Now the structure of the stamen is gradually differentiated, with locules developing on the anthers (Figure 5E, F). The anthers continue to develop, and the pollen matures at stage M6 (Figure 5G, H). The male flower is fully mature at stage M7. The sepals open fully, allowing the anther to open, and pollen is released (Figure 5J, K). The flowers wither and detach from the pedicel at stage M8 (Figure 5L, M). Importantly, there was no evidence of the formation or even initiation of a carpel whorl at any stage of hemp male flower development.

**Figure 5.**
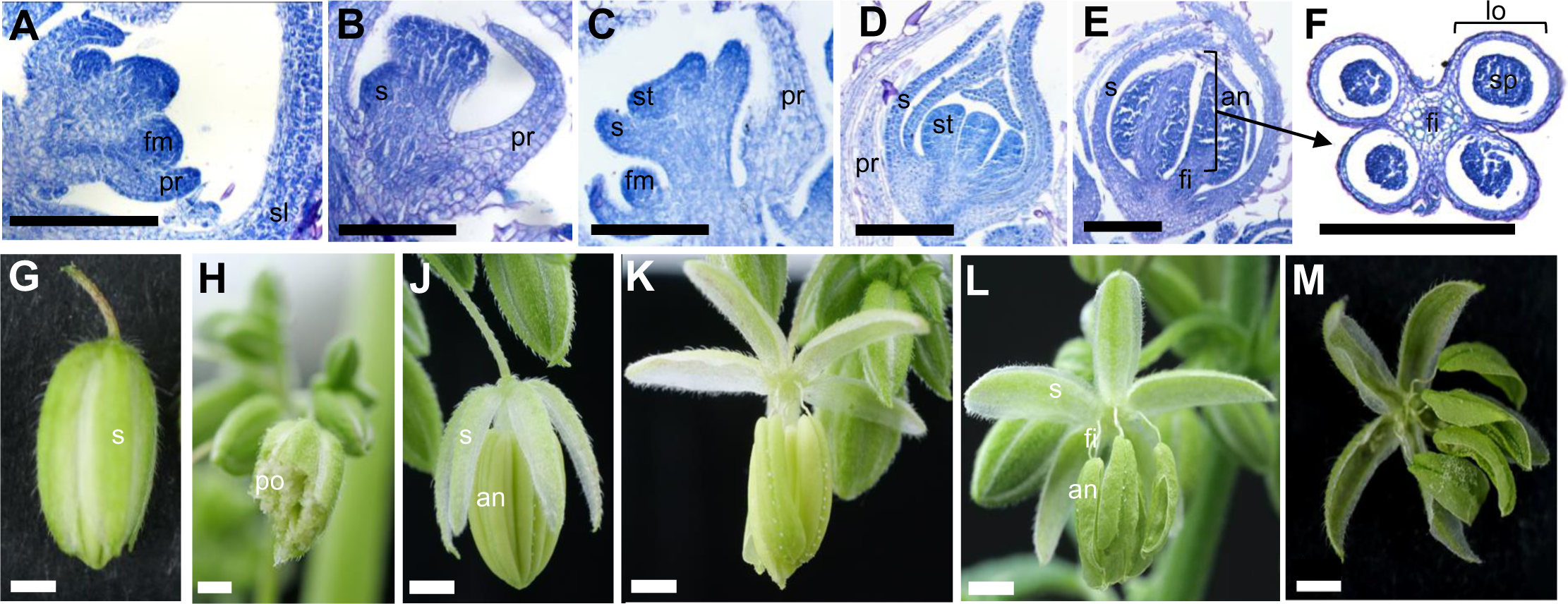
Male flower development in *C. sativa*. Sections display male hemp flower development. At stage M1, the floral meristem is visible (A). At stage M2, the sepal primordium is visible (B). At stage M3, stamen primordia have formed (C). At stage M4, sepals have enclosed the developing stamen (D). At stage M5, anthers differentiate as shown in longitudinal (E) and transverse sections (F). Late flowering stages are imaged using macrophotography. At stage M6, pollen continues developing (G, H). At stage M7, the flower is at anthesis (J, K). At stage M8, flowers undergo senescence (L, M). an. Anther; fi. Filament; fm. Floral meristem; lo. Locule; po. Pollen; pr. Prophyll; s. Sepal or sepal primordium; sl. Subtending leaf; sp. Sporogenous tissue; st. Stamen or stamen primordium. Scale bar: (A) 200 µm; (B to F) 100 µm; (G to M) 1 mm.

In female hemp plants, flowering stage 1 (stage F1) is characterized by the emergence of a floral meristem from the flanks of the inflorescence, with a developing prophyll extending under it (Figure 6A). In stage F2, a bract primordium develops on one side of the floral meristem (Figure 6B). In stage F3, a carpel primordium emerges, and at the base of the carpel, the perianth develops (Figure 6C). At stage F4, the bract extends and the perianth is well developed and surrounds the lower part of the elongated carpel primordia (Figure 6D). In stage F5, two stigmas differentiate and elongate from the carpel primordia (Figure 6E). During stage F6, the ovary is developing (Figure 6F). The stigma is fully extended at stage F7 and the ovule is developed (Figure 6G, H, J). The final stage F8 of female flower development is the initiation of seed development if pollination is successful. The ovary keeps developing, and the stigma turns red and wither (Figure 6K, L). No stamen whorl was observed at any stage of development in female hemp flowers.

**Figure 6.**
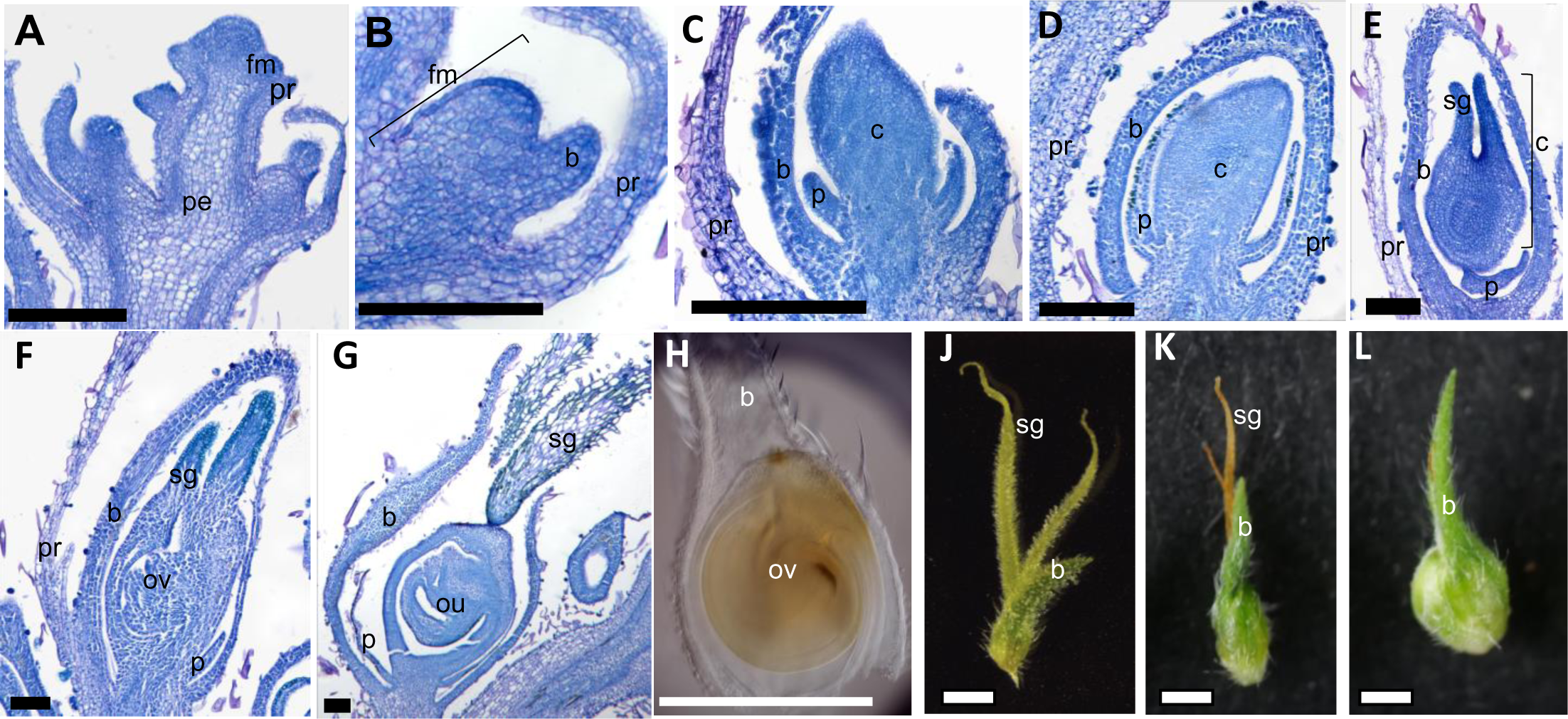
Female flower development in *C. sativa*. Sections of developing female hemp flowers show meristem and flower at early flowering stages. At stage F1, the floral meristem is established (A). At stage F2, a bract primordium is established (B). At stage F3, the perianth and carpel begin developing (C). At stage F4, the bract and perianth are extended and the carpel continues development (D). At stage F5, stigmata differentiate (E). At stage F6, the ovary is developing (F). At stage F7, the stigmata are fully extended and the ovule is clearly developed as shown in section (G), the image from a stereomicroscope (H) and macrophotography (J). Following fertilisation, the seed develops and stigmata senesce (K, L). b. Bract; c. Carpel or carpel primordium; fm. Floral meristem; ou. Ovule; ov. Ovary; p. Perianth; pe. Peduncle; pr. Prophyll; sg. Stigma. Scale bar: (A to G) 100 µm; (H to L) 1 mm.

### Expression of putative floral regulators

Male and female hemp plants of the cultivar ‘FINOLA’ could be macroscopically distinguished at the fourth true leaf stage L4 (Figure 2). This indicated that gene expression changes that govern male vs. female hemp development takes place at an even earlier developmental stage. To study this in more detail, we analysed the expression of the three putative key regulatory genes of flower development.

In Arabidopsis, *SOC1*, *LFY* and *AG* are among the key regulatory genes controlling flower development (Lee and Lee, 2010; Ó’Maoiléidigh et al., 2013). The roles of those genes are broadly conserved throughout flowering plant evolution, they therefore serve as good indicators of a switch from vegetative to reproductive development (Chávez-Hernández et al., 2022). *CsSOC1* and *CsAG* have been identified before (Wan et al., 2021). We identified the *C. sativa* genes *CsLFY* as putative orthologs of *LFY* from Arabidopsis (Figure S1). Expression was studied in shoot apices starting from the second true leaf stage (stage L2). The expression of these three genes in mature flowers (stage M7 and stage F7) and vegetative leaves serve as additional comparators.

Expression of all three *C. sativa* genes was analysed via RT-qPCR. The results indicate that the expression of *CsSOC1* does not change significantly in the apical meristems of male plants from stage L2 to stage L4 (Figure 7A). In contrast, *CsSOC1* has a higher expression in female hemp plants at L8 just before the onset of flowering as compared to early vegetative stages L2 to L4 (Figure 7B).

**Figure 7.**
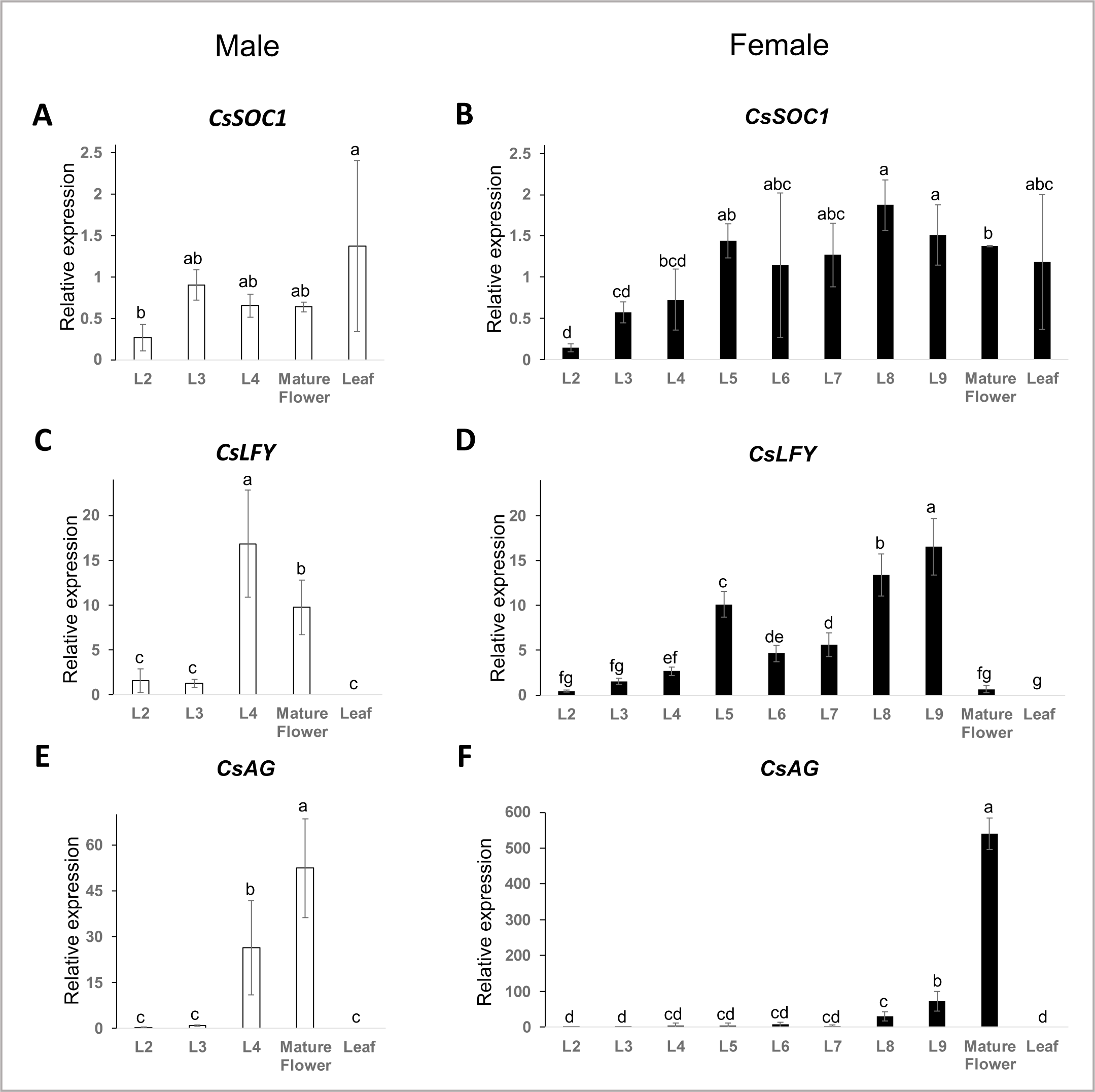
Expression of genes putatively involved in flower initiation and development in *C. sativa*. Expression analysis using RT-qPCR of *CsSOC1* (A, B), *CsLFY* (C, D) and *CsAG* (E, F). Expression was measured in male (A, C, E) and female (B, D, F) apical shoot meristem of different developmental stages (L2 to L9), mature flowers (stage M7 and F7) and vegetative leaves. The expression value of the genes was performed in 2-delta delta Ct value (2-ΔΔCt). The comparative Ct (ΔΔCt) values were directly produced from QuantStudio 6 and 7 Pro RealTime PCR Systems Software (Thermo Fisher Scientific, America) using housekeeping gene *UBQ* and *PP2A* for endogenous control and computed as the geometric mean. Biological replicates (n=3) for each stage were included. Error bars indicate standard deviations. The Tukey test (Honestly significant difference test) was used to assess the significance of differences between groups. Significance levels are indicated by letters.

The expression of *CsLFY* increased relatively strongly from stage L3 to stage L4 in male hemp plants (Figure 7 C). Likewise, the highest expression of *CsLFY* in female plants was observed at the start of flowering at L9 (Figure 7 D). Interestingly, it appears the highest expression of *CsLFY* in both male and female plants is at the previously identified flowering stage (Figure 7C, D).

The putative floral organ identity gene *CsAG* maintained a low level of expression in male hemp apical meristems at early growth stages L2 and L3. *CsAG* expression increased strongly at stage L4 when male flowers became visible macroscopically. The expression of *CsAG* was further increased in mature male hemp flowers (Figure 7E). The expression level of the *CsAG* gene in female hemp plants was significantly increased at stage L9. Likewise, the highest expression level of CsAG was observed in mature female hemp flowers (Figure 7F).

## Discussion

Previous studies on flower development in dioecious plants have identified two major groups based on the mode of sex organ development (Heslop-Harrison, 1957; Matsunaga and Kawano, 2001). In the first group, the flowers of dioecious plants start to develop as hermaphrodites with both sex organ types, but male or female organs are aborted or suppressed during development, and the plant subsequently becomes unisexual. For instance, in the white campion (*Silene latifolia*), in male flowers the development of the carpel primordium is initiated, but the carpel remains rudimentary in the later stages of male flower development. Likewise, stamens start to develop in the female flowers but later undergo degeneration as the carpel matures (Grant et al., 1994). The sexual organs of the opposite sex may even develop to a more advanced stage of flower development in some other dioecious plants like asparagus (*Asparagus officinalis*) (Caporali et al., 1994)

In contrast to the first group, the opposite reproductive organ set (carpels in the male and stamens in the female) in the second group of plants is not initiated at all. This is the case in plants like hop (*Humulus lupulus*) and spinach (*Spinacia oleracea*) (Shephard *et al*., 2000; Sherry, Eckard and Lord, 1993). Our study confirms earlier reports (Leme et al., 2020) that hemp belongs to the second group of plants: male flowers do not show any signs of carpel initiation, while female flowers develop no stamens. However, the differences between male and female flowers extend substantially beyond the presence/absence of carpels and stamens: the structure of male and female flowers is different, with male flowers possessing five prominent sepals whereas female flowers develop a perianth that is hardly visible macroscopically. Differences also affect flowering time, with male plants flowering earlier than female plants. It should be noted, however, that while female flowers emerge later, once they become visible, their stigmata emerge and flowers and ovules are developed, ready to receive pollen. In contrast, male *C. sativa* flowers undergo significant further development after first being visible macroscopically. Also, the inflorescence structure between male and female plants is different: whereas female ‘FINOLA’ hemp plants develop a spike, male plants develop a panicle under the growth conditions applied.

The difference in flowering time is also present at the molecular level. The expression of the putative floral integrator *CsSOC1*: In male hemp plants, the expression of *CsSOC1* remained relatively consistent throughout their growth stages. In contrast, a distinct elevation in *CsSOC1* expression was observed in female hemp plants, particularly at stages L8 and L9, when compared to the early vegetative stages (L2 to L4). The combination of molecular and morphological data points to significant variations between male and female *C. sativa* plants.

Together, morphological and molecular data indicate profound differences between male and female *C. sativa* plants. In many plant species for which sex determining genes have been identified those genes act at the level of pollen or carpel development, i.e. relatively late during development (Leite Montalvão et al., 2021). We speculate that any sex determining gene(s) in *C. sativa* will act much earlier during development, also contributing to differences in flowering time and inflorescence architecture. Later flowering in female photoperiod-insensitive *C. sativa* individuals might be necessary or advantageous, as a longer vegetative phase might give female plants more time to develop foliage important to sustain developing seeds.

It is noteworthy that other species within the Cannabaceae are also dioecious, but stamens or carpels are aborted much later during development than in *C. sativa* (Leme et al., 2020). It will be interesting to study the developmental control and evolutionary origin of this morphological variation in more detail.

### Landmark developmental events in *Cannabis sativa*

We observe a relatively robust initiation of male flower development at the L3/L4 stage, whereas female flowers appear later, at around stage L8/L9. At least in our experiments, greater variability was observed when expressing flowering time in days after sowing instead of the number of leaf pairs. Similar observations are made in Arabidopsis (Karlsson et al., 1993), where flowering time is now routinely estimated as the number of rosette leaves at the time of flowering. The cultivar ‘FINOLA’ studied here is photoperiod insensitive, which means the flowering time is independent of day length (Schilling et al., 2023). Days until flowering and number of leaf pairs developed until flowering will inevitably depend on the day length in photoperiod sensitive cultivars. It will therefore be interesting to explore how robust leaf development is as a proxy for flowering time in other cultivars and environmental conditions. The order of events in flower development in *C. sativa* is broadly similar to other eudicots (Bowman et al., 1991; Becker et al., 2005), with sepals emerging before the reproductive organs. It is interesting that a perianth is clearly identifiable in young female flowers, but withers at later stages of development, with a bract enclosing the carpel of the mature female flower (Leme et al., 2020). The perianth is more pronounced in male flowers, which form five robust sepals. The bract of the female flowers but not the sepals of the male flowers develop glandular trichomes producing cannabinoids (Livingston et al., 2020). This underscores the profound morphological difference between male and female flowers and indicates that the sex determination mechanism may activate distinct gene regulatory circuits in male and female flowers. The stages of flower development specified here will be helpful in further finding the genes and gene regulatory networks involved in male vs. female flower development.

### The expression of candidate flowering-related genes is similar in male and female hemp plants

The expression of *CsSOC1* did not significantly vary in male hemp plants in the developmental stages analysed. On the other hand, in female hemp plants, there was a noticeable increase in expression at stages L8 and L9 as compared to early vegetative stages L2 to L4. This may reflect differences in flowering time between male and female ‘FINOLA’ hemp plants: male plants flower extremely early, so *CsSOC1* expression might be high in very early development already, whereas female plants flower later. *CsLFY* showed the highest expression level at stage L4 in males and stage L9 in females, when the apical inflorescence was visible and the transition from the vegetative stage to the reproductive stage occurred. Furthermore, the expression of *CsAG* reaches its maximum in mature hemp flowers.

Overall, the expression of the *C. sativa* genes is remarkably similar to their Arabidopsis homologs: *SOC1* in Arabidopsis is a flowering time gene and expressed during the vegetative stages before flowering (Liu et al., 2009). *LFY* activates the floral homeotic genes that determine floral organ identity (Blazquez et al., 1997; Liu et al., 2009). The expression of *LFY* in Arabidopsis increases when the meristem transitions from a vegetative to an inflorescence meristem (Schmid et al., 2005). *AG* specifies carpel and stamen identity and shows high expression levels in those organs (Okamuro et al., 1993; Schmid et al., 2005). This conservation in expression between *C. sativa* and Arabidopsis indicates functional conservation of the genes. The observation that all genes analysed here are upregulated in male as well as female flowers further indicates that core pathways regulating flower development are similar in male and female flowers. However, upstream regulators may express differently in the molecular mechanisms of male and female plants.

In conclusion, this study provided a comprehensive analysis of hemp male and female inflorescence and flower development. By identifying landmark events and highlighting the early stage divergence between male and female plants, this study provides insights into the reproductive development of hemp. The differential expression of key marker genes at different time points further showed the complexities of sex-specific development in male and female hemp plants at the molecular level.

## Acknowledgement

JS is supported by a Chinese Research Council Postgraduate Scholarship (Grant No.: CSC No. 201908300031).

## Supplementary Material

### Supplementary Tables

**Table S1.**
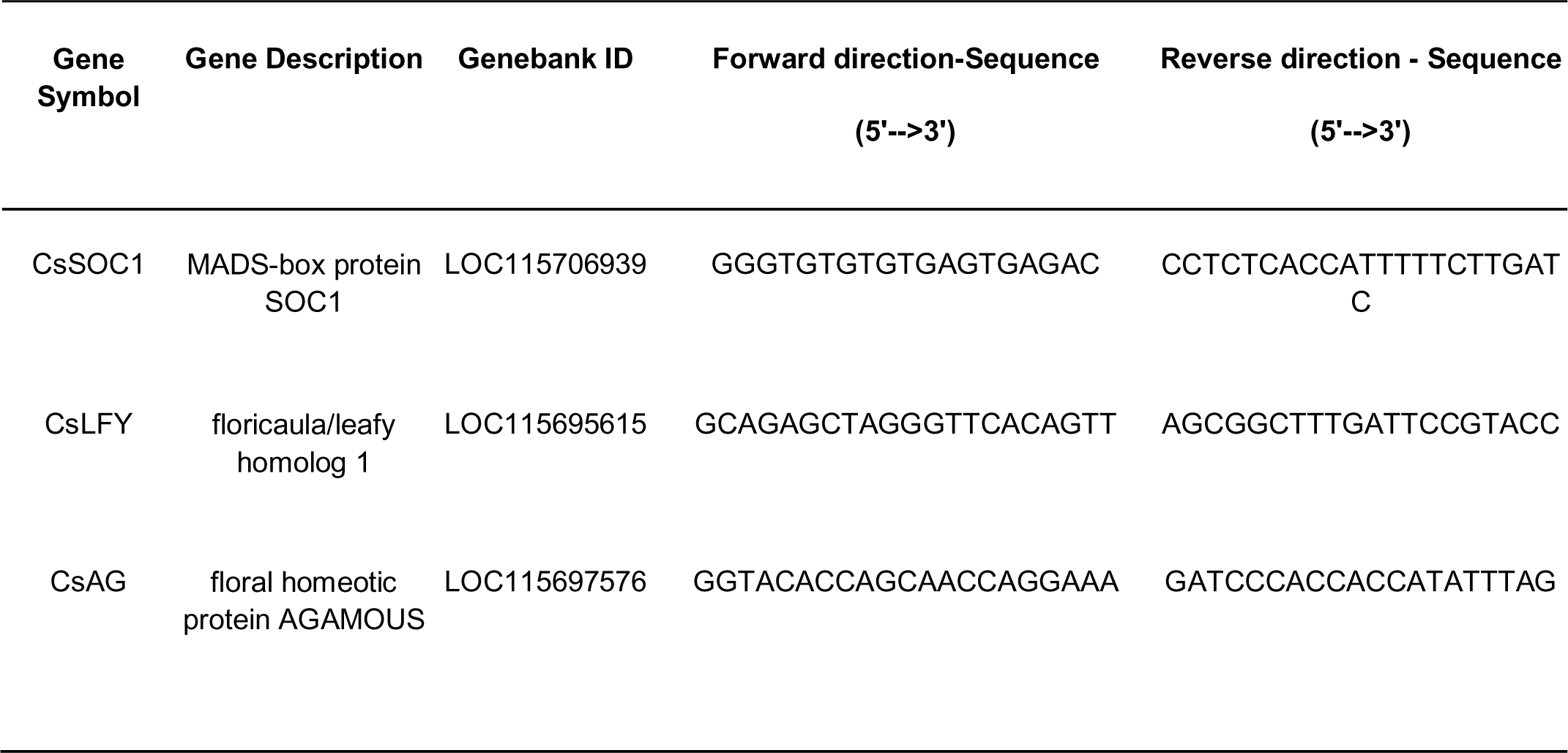
Three candidate genes and their primer sequences for RT-qPCR.

### Supplementary figures

**Figure S1.**
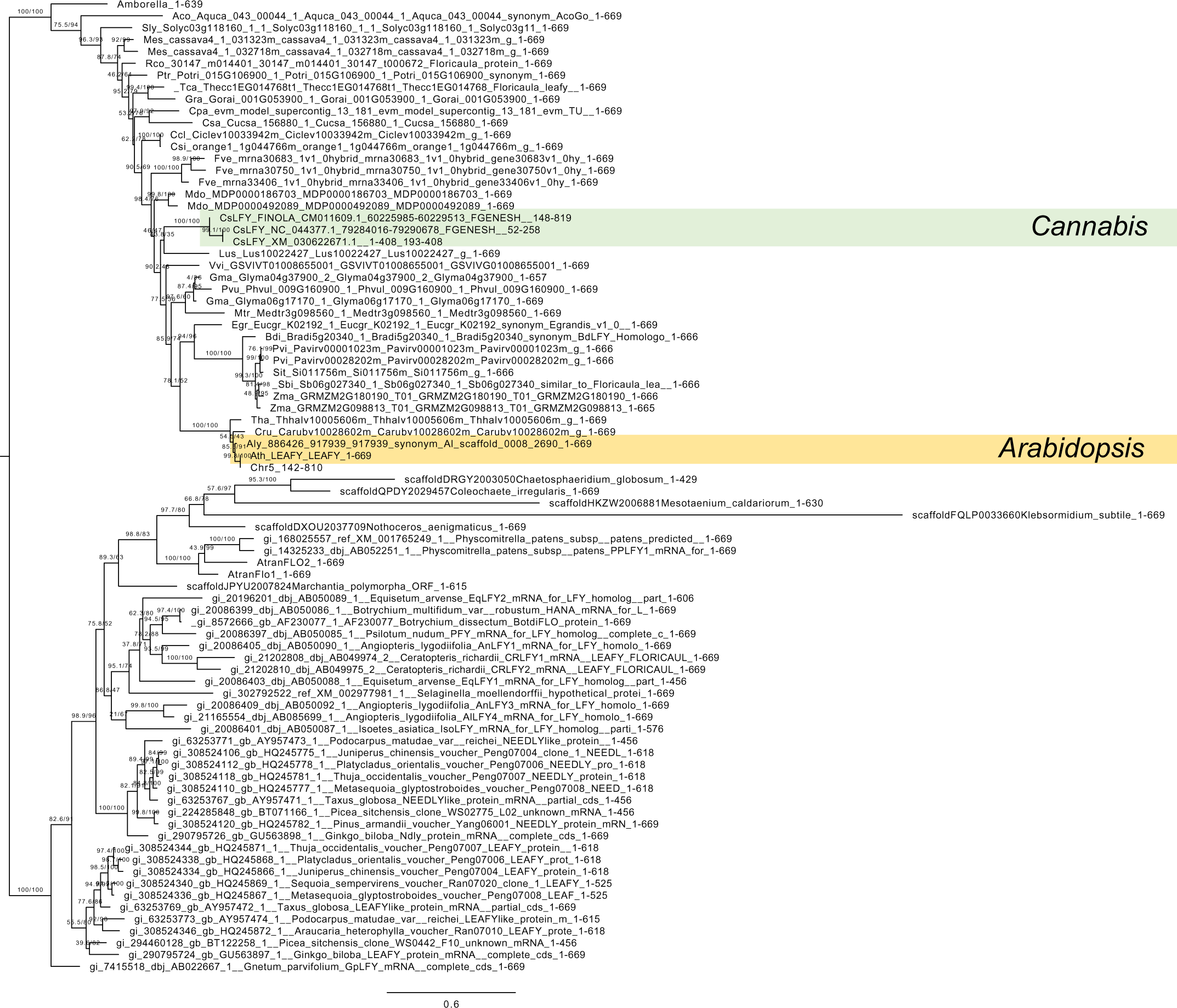
*CsLFY* is closely related to *LFY* from Arabidopsis. A maximum likelihood tree was generated using gymnosperm and angiosperm *LFY* orthologs previously published (Sayou et al., 2014) as well as *CsLFY* identified in *C. sativa. CsLFY* (green background) was identified through BLAST (Altschul et al., 1990) against the *C. sativa* reference genome (GCA_900626175.2; LOC115695615; XM_030622671.1) (Grassa et al., 2021). Additionally, gene predictions for *CsLFY* were made using FGenesH+ (Solovyev, 2004), LFY (AT5G61850, orange background) and the genomic region of the reference genome (NC_044377.1:79284016-79290678) as well as the hemp cultivar ‘FINOLA’ (GCA_003417725.2; CM011609.1:60225985-60229513). All coding sequences were aligned using MAFFT (Katoh et al., 2005) and trimmed to include sections homologous to *CsLFY*. The maximum likelihood tree was generated using IQtree (Trifinopoulos et al., 2016) and visualized using FigTree (http://tree.bio.ed.ac.uk/software/figtree/).

## References

Aizpurua-Olaizola, O., Soydaner, U., Öztürk, E., Schibano, D., Simsir, Y., Navarro, P., Etxebarria, N., Usobiaga, A., 2016. Evolution of the Cannabinoid and Terpene Content during the Growth of Cannabis sativa Plants from Different Chemotypes. J. Nat. Prod. 79, 324–331. 10.1021/acs.jnatprod.5b00949

Altschul, S.F., Gish, W., Miller, W., Myers, E.W., Lipman, D.J., 1990. Basic local alignment search tool. J Mol Biol 215, 403–410. 10.1016/S0022-2836(05)80360-2

Blazquez, M.A., Soowal, L.N., Lee, I., Weigel, D., 1997. LEAFY expression and flower initiation in Arabidopsis. Development 124, 3835–3844. 10.1242/dev.124.19.3835

Bowman, J.L., Smyth, D.R., Meyerowitz, E.M., 1991. Genetic interactions among floral homeotic genes of Arabidopsis. Development 112, 1–20. 10.1242/dev.112.1.1

Caporali, E., Carboni, A., Galli, M.G., Rossi, G., Spada, A., Marziani Longo, G.P., 1994. Development of male and female flower in Asparagus officinalis. Search for point of transition from hermaphroditic to unisexual developmental pathway. Sexual Plant Reprod 7, 239–249. 10.1007/BF00232743

Chávez-Hernández, E.C., Quiroz, S., García-Ponce, B., Álvarez-Buylla, E.R., 2022. The flowering transition pathways converge into a complex gene regulatory network that underlies the phase changes of the shoot apical meristem in Arabidopsis thaliana. Frontiers in Plant Science 13.

Feng, G., Sanderson, B.J., Keefover-Ring, K., Liu, J., Ma, T., Yin, T., Smart, L.B., DiFazio, S.P., Olson, M.S., 2020. Pathways to sex determination in plants: how many roads lead to Rome? Current Opinion in Plant Biology 54, 61–68. 10.1016/j.pbi.2020.01.004

Finnan, J., Burke, B., 2013. Nitrogen fertilization to optimize the greenhouse gas balance of hemp crops grown for biomass. GCB Bioenergy 5, 701–712. 10.1111/gcbb.12045

Finnan, J., Styles, D., 2013. Hemp: A more sustainable annual energy crop for climate and energy policy. Energy Policy 58, 152–162. 10.1016/j.enpol.2013.02.046

Fraguas-Sánchez, A.I., Torres-Suárez, A.I., 2018. Medical Use of Cannabinoids. Drugs 78, 1665–1703. 10.1007/s40265-018-0996-1

Grant, S., Hunkirchen, B., Saedler, H., 1994. Developmental differences between male and female flowers in the dioecious plant Silene latifolia. The Plant Journal 6, 471–480. 10.1046/j.1365-313X.1994.6040471.x

Grassa, C.J., Weiblen, G.D., Wenger, J.P., Dabney, C., Poplawski, S.G., Timothy Motley, S., Michael, T.P., Schwartz, C.J., 2021. A new Cannabis genome assembly associates elevated cannabidiol (CBD) with hemp introgressed into marijuana. New Phytologist 230, 1665–1679. 10.1111/nph.17243

Heslop-Harrison, J., 1957. The Experimental Modification of Sex Expression in Flowering Plants. Biological Reviews 32, 38–90. 10.1111/j.1469-185X.1957.tb01576.x

Karlsson, B.H., Sills, G.R., Nienhuis, J., 1993. Effects of Photoperiod and Vernalization on the Number of Leaves at Flowering in 32 Arabidopsis thaliana (Brassicaceae) Ecotypes. American Journal of Botany 80, 646–648. 10.2307/2445435

Katoh, K., Kuma, K., Toh, H., Miyata, T., 2005. MAFFT version 5: improvement in accuracy of multiple sequence alignment. Nucleic Acids Res 33, 511–518. 10.1093/nar/gki198

Lee, J., Lee, I., 2010. Regulation and function of SOC1, a flowering pathway integrator. Journal of Experimental Botany 61, 2247–2254. 10.1093/jxb/erq098

Leite Montalvão, A.P., Kersten, B., Fladung, M., Müller, N.A., 2021. The Diversity and Dynamics of Sex Determination in Dioecious Plants. Frontiers in Plant Science 11.

Leizer, C., Ribnicky, D., Poulev, A., Dushenkov, S., Raskin, I., 2000. The Composition of Hemp Seed Oil and Its Potential as an Important Source of Nutrition. Journal of Nutraceuticals, Functional & Medical Foods 2, 35–53. 10.1300/J133v02n04_04

Leme, F.M., Schönenberger, J., Staedler, Y.M., Teixeira, S.P., 2020. Comparative floral development reveals novel aspects of structure and diversity of flowers in Cannabaceae. Botanical Journal of the Linnean Society 193, 64–83. 10.1093/botlinnean/boaa004

Liu, C., Xi, W., Shen, L., Tan, C., Yu, H., 2009. Regulation of Floral Patterning by Flowering Time Genes. Developmental Cell 16, 711–722. 10.1016/j.devcel.2009.03.011

Livingston, S.J., Quilichini, T.D., Booth, J.K., Wong, D.C.J., Rensing, K.H., Laflamme-Yonkman, J., Castellarin, S.D., Bohlmann, J., Page, J.E., Samuels, A.L., 2020. Cannabis glandular trichomes alter morphology and metabolite content during flower maturation. The Plant Journal 101, 37–56. 10.1111/tpj.14516

Matsunaga, S., Kawano, S., 2001. Sex Determination by Sex Chromosomes in Dioecious Plants. Plant Biol (Stuttg) 3, 481–488. 10.1055/s-2001-17735

McMaster, G.S., Moragues, M., 2019. Crop Development Related to Temperature and Photoperiod, in: Savin, R., Slafer, G.A. (Eds.), Crop Science, Encyclopedia of Sustainability Science and Technology Series. Springer, New York, NY, pp. 9–28. 10.1007/978-1-4939-8621-7_384

Okamuro, J.K., den Boer, B.G., Jofuku, K.D., 1993. Regulation of Arabidopsis flower development. Plant Cell 5, 1183–1193.

Ó’Maoiléidigh, D.S., Wuest, S.E., Rae, L., Raganelli, A., Ryan, P.T., Kwaśniewska, K., Das, P., Lohan, A.J., Loftus, B., Graciet, E., Wellmer, F., 2013. Control of Reproductive Floral Organ Identity Specification in Arabidopsis by the C Function Regulator AGAMOUS. The Plant Cell 25, 2482–2503. 10.1105/tpc.113.113209

Ryan, P.T., Ó’Maoiléidigh, D.S., Drost, H.-G., Kwaśniewska, K., Gabel, A., Grosse, I., Graciet, E., Quint, M., Wellmer, F., 2015. Patterns of gene expression during Arabidopsis flower development from the time of initiation to maturation. BMC Genomics 16, 488. 10.1186/s12864-015-1699-6

Sayou, C., Monniaux, M., Nanao, M.H., Moyroud, E., Brockington, S.F., Thévenon, E., Chahtane, H., Warthmann, N., Melkonian, M., Zhang, Y., Wong, G.K.-S., Weigel, D., Parcy, F., Dumas, R., 2014. A promiscuous intermediate underlies the evolution of LEAFY DNA binding specificity. Science 343, 645–648. 10.1126/science.1248229

Schilling, S., Melzer, R., Dowling, C.A., Shi, J., Muldoon, S., McCabe, P.F., 2023. A protocol for rapid generation cycling (speed breeding) of hemp (Cannabis sativa) for research and agriculture. The Plant Journal 113, 437–445. 10.1111/tpj.16051

Schmid, M., Davison, T.S., Henz, S.R., Pape, U.J., Demar, M., Vingron, M., Schölkopf, B., Weigel, D., Lohmann, J.U., 2005. A gene expression map of Arabidopsis thaliana development. Nat Genet 37, 501–506. 10.1038/ng1543

Shephard, H.L., Parker, J.S., Darby, P., Ainsworth, C.C., 2000. Sexual development and sex chromosomes in hop. New Phytologist 148, 397–411. 10.1046/j.1469-8137.2000.00771.x

Sherry, R.A., Eckard, K.J., Lord, E.M., 1993. Flower Development in Dioecious Spinacia oleracea (Chenopodiaceae). American Journal of Botany 80, 283–291. 10.2307/2445351

Silvestro, S., Mammana, S., Cavalli, E., Bramanti, P., Mazzon, E., 2019. Use of Cannabidiol in the Treatment of Epilepsy: Efficacy and Security in Clinical Trials. Molecules 24, 1459. 10.3390/molecules24081459

Smyth, D.R., Bowman, J.L., Meyerowitz, E.M., 1990. Early flower development in Arabidopsis. The Plant Cell 2, 755–767. 10.1105/tpc.2.8.755

Solovyev, V., 2004. Statistical Approaches in Eukaryotic Gene Prediction, in: Handbook of Statistical Genetics. 10.1002/0470022620.bbc06

Spitzer-Rimon, B., Duchin, S., Bernstein, N., Kamenetsky, R., 2019. Architecture and Florogenesis in Female Cannabis sativa Plants. Front. Plant Sci. 10, 350. 10.3389/fpls.2019.00350

Trifinopoulos, J., Nguyen, L.-T., von Haeseler, A., Minh, B.Q., 2016. W-IQ-TREE: a fast online phylogenetic tool for maximum likelihood analysis. Nucleic Acids Research 44, W232–W235. 10.1093/nar/gkw256

Wan, Z., Lu, M., Wu, S., Mi Y., Zhai, J., 2021. Identification and expression analysis of the MIKC-type MADS-box gene family in Cannabis sativa L.[J]. Acta Pharmaceutica Sinica, 2021,56(11): 3173-3183. 10.16438/j.0513-4870.2021-0892

